# *Pseudomonas aeruginosa* displays a dormancy phenotype during long-term survival in water

**DOI:** 10.1101/327163

**Authors:** Shawn Lewenza, Jason Abboud, Karen Poon, Madison Kobryn, Istvan Humplik, John Rainer Bell, Laura Mardan, Shauna Reckseidler-Zenteno

**Affiliations:** Faculty of Science and Technology, Athabasca University, 1 University Drive, Athabasca, Alberta, Canada T9S 3A3; Department of Microbiology, Immunology, and Infectious Diseases, Faculty of Medicine, University of Calgary, 3330 Hospital Drive NW, Calgary, Alberta, Canada T2N 4N1

**Keywords:** dormancy, *Pseudomonas aeruginosa*, survival, low nutrient environments, water, metabolism, membrane, nosocomial pathogen, gene expression, persistence, reservoir

## Abstract

*Pseudomonas aeruginosa* is capable of long-term survival in water, which may serve as a reservoir for infection. Although viable cell counts of PAO1 incubated in water remain stable throughout 8 weeks, LIVE/DEAD® staining indicated a high proportion of cells stained with propidium iodide (PI). The proportion of PI-stained cells increased by 4 weeks, then decreased again by 8 weeks, suggesting an adaptive response. This was also evident in an observed shift in cell morphology from a rod to a coccoid shape after 8 weeks. Fluorescence-activated cell sorting (FACS) was used to recover PI-stained cells, which were plated and shown to be viable, indicating that PI-stained cells were membrane-compromised but still cultivable. PAO1 mid-log cells in water were labeled with the dsDNA-binding dye PicoGreen® to monitor viability as well as DNA integrity, which demonstrated that the population remains viable and transitions towards increased dsDNA staining. Metabolic activity was found to decrease significantly in water by 4 weeks. The PAO1 outer membrane became less permeable and more resistant to polymyxin B damage in water, and the profile of total membrane lipids changed over time. None of the individual mutants within a library of ~2500 mapped, mini-Tn*5-lux* transposon mutants were found to have decreased survival in water. Among the ~1400 transcriptional *lux* fusions, gene expression in water revealed that the majority of genes were repressed, but subsets of genes were induced at particular time points. In summary, these results indicate that *P. aeruginosa* is dormant in water and this adaptation involves a complex pattern of gene regulation and changes to the cell to promote long-term survival and antibiotic tolerance. The approach of *P. aeruginosa* incubated in water may be useful to study antibiotic tolerance and the mechanisms of dormancy and survival in nutrient limiting conditions.

## INTRODUCTION

*Pseudomonas aeruginosa* is a Gram-negative bacterium that is ubiquitous in the environment and appreciated for its ability to cause disease in plants, insects, animals, and humans [1,2]. This opportunistic organism is a major cause of nosocomial or hospital-acquired infections, most notably causing skin infections in burn patients and diabetic foot wounds, infections of indwelling devices such as catheters, and chronic lung infections in patients with Cystic Fibrosis [3]. Studies have shown that *P. aeruginosa* may survive for months on hospital surfaces [4]. *P. aeruginosa* is an archetypal biofilm-forming organism, which is a conserved strategy used for long-term survival in nature and during infections. *P. aeruginosa* infections are very difficult to treat because the organism has numerous intrinsic antibiotic resistance mechanisms, and grows as multidrug tolerant biofilms, thus promoting chronic infections [3].

*P. aeruginosa* has been shown to survive in water for over 145 days (20.7 weeks), significantly longer than two other bacterial pathogens, *Escherichia coli* and *Staphylococcus aureus* [5]. The ability of the organism to persist at length in water indicates that water may be an environmental reservoir for *P. aeruginosa*. The microbiome of pristine groundwater is dominated by the *Pseudomonas* genus, which was shown to represent 10% of all species [6]. Numerous studies have demonstrated that *P. aeruginosa* can be isolated from hospitals, from the water in intensive care units, as well as sinks, basins, drains, showers, toilets and bathtubs, leading to transmission of *P. aeruginosa* infections [7,8]. It is therefore important to understand how this organism is able to survive in water, to better understand the transmission and possibly improve infection control policies.

Slow or non-growing states are poorly understood, yet bacteria in nature exist most commonly in nutrient limited conditions. There are numerous experimental systems used to study nongrowing bacteria, which include persister cells, starved cells in stationary phase, or the viable-but-nonculturable (VBNC) state [9]. All of these non-growing states can be considered a form of dormant bacterial cells. Non-growing persister cells are present in laboratory grown planktonic and biofilm cultures, and contribute to multidrug antibiotic tolerance and chronic infections [10]. Persister cells are not utilizing nutrients, producing proteins, synthesizing any replication machinery, and therefore not multiplying [10,11]. However, starved cells in stationary phase have also been studied as a model of non-growing cells and were shown to maintain constant gene expression and protein production during extended starvation periods [12]. Both persister cells and starved stationary phase cells are dormant growth states that are capable of growth once they are reintroduced into a nutrient-rich conditions [10,12]. The objective of this study was to characterize the long-term survival of *P. aeruginosa* in water and to determine if this experimental system is a useful dormancy model for future studies of bacterial survival in nutrient limiting conditions.

## MATERIALS AND METHODS

### Bacterial strains used in this study

*P. aeruginosa* PAO1 was utilized as the wild type strain in this study. PAO1 was grown in Luria broth (LB) and incubated at 37°C with shaking at 250 rpm overnight. Stationary phase cultures (ON) were obtained following overnight growth in LB and these were inoculated into water. Logarithmic phase cultures (log) were obtained by sub-culturing overnight cultures into LB and growing these to an optical density (OD_600_) of 0.5 before being inoculated into water for experimental procedures. Samples were removed from water at a number of time points for analysis.

### Preparation of water samples and quantitative determination of *P. aeruginosa* viability in water

Strains were grown overnight or to mid-log in 3mL LB and an aliquot of 1 ml of the cultures was centrifuged at 13,000 rpm for 3 min., washed three times in sterile distilled H_2_O (sdH_2_O), and resuspended in a final volume of 1 ml sdH_2_O. The washed culture was used to inoculate sdH_2_O at a final concentration of 10^7^ CFU/ml. The tubes were loosely capped and incubated at room temperature. Bacterial quantitation of the samples was performed to accurately determine the CFU/ml at time zero and at various time points in water.

### Live/dead staining and fluorescence-activated cell sorting (FACS) of *P. aeruginosa* strains incubated in water

At each time point, 1 ml of *P. aeruginosa* in water (10^7^ CFU) was removed, centrifuged at 13,000 rpm for 3 min and resuspended in 1 ml of 0.9% saline. Samples were then subjected to LIVE/DEAD® staining using the BacLight™ kit comprised of SYTO 9 (green) and propidium iodide (PI; red) (Thermo Fisher), which were added at a final concentration of 30 and 10 μM, respectively. Samples were then subjected to flow cytometry to quantitatively determine the proportion of SYTO9 and PI-stained cells. Once the proportion of green/red cells was determined, the green and red cell sub-populations were separated into fractions by fluorescence-activated cell sorting (FACS). The viability of cells in each fraction was determined by serially diluting and plating of the sorted fractions onto LB agar. The Quant-iT™ PicoGreen® dsDNA reagent was used to quantitate dsDNA (1 μl of the stock solution was added to 250 μl of bacteria sample in water) (Thermo Fisher). A total of 50,000 events were acquired for all flow cytometry experiments in list mode files and analyzed using BD FACS Diva software.

### Fluorescence microscopy of live/dead stained cells

At each time point, 1 ml of the water sample containing PAO1 was centrifuged and resuspended in 0.9% saline. The LIVE/DEAD® reagents were added to 10 μl of the sample at a concentration of 30 and 10 μM respectively and 2 μl of the sample was added to an agarose bed-coated glass slide and sealed with a glass coverslip. Slides were visualized on a Leica DMI 4000 B wide field fluorescence microscope.

### Assessment of ATP production of PAO1 in water

PAO1 incubated in water was assessed over time for metabolic activity based on ATP production. ATP production was determined using the BacTiter-Glo™ Microbial Cell Viability Assay (Promega), which measures the amount of ATP present in a sample as a function of luminescence. The BacTiter-Glo™ reagent was added to the PAO1 water sample at a ratio of 1:1 and luminescence was measured in counts per second after 5 min incubation at room temperature.

### NPN assay to measure outer membrane (OM) permeability

OM permeability of PAO1 was assessed using a previously established protocol [13,14]. Samples of PAO1 in LB or water (1 ml, 10^7^ CFU) were centrifuged at 13, 000 rpm for 2 min and cells were resuspended in 5 mM HEPES buffer (pH 7.2) containing 5 mM glucose. Cells were pretreated with sodium azide (0.2%) to disable active efflux. 1-N-phenylnaphthylamine (NPN) is a fluorescent dye when integrated into the hydrophobic environment of bacterial membranes. NPN was added to measure both the baseline permeability and to measure the outer membrane tolerance to polymyxin B treatment. After NPN addition, cells were treated with polymyxin B (5 μg/ml) to disrupt the OM, leading to increased NPN uptake and fluorescence, which was measured using a Spectra Max M2 spectrophotometer using the SoftMax Pro 6 software. The setting for green excitation and emission spectra were set to 350nm and 420nm respectively, at 30°C. Samples of PAO1 were prepared in triplicate for each assay, and the assay was performed three times.

### Lipid detection of PAO1 cells following incubation in water

The total lipid profile of PAO1 cells following incubation in water was determined by thin-layer chromatography as previously described [15]. Total lipids were extracted with chloroform:methanol (1:2), separated using thin layer chromatography and amino group containing lipids were visualized by spraying with the ninhydrin reagent (Sigma). Lipids were isolated from PAO1 incubated in water, and from control PAO1 cultures grown in low and high phosphate BM2-defined growth media [15].

### Gene expression and survival in water analysis

All mapped transposon mutants (containing about 50% transcriptional *lux* fusions) from a previously constructed transposon mutant library [16] were re-arrayed into a series of 96-well microplates. The mutants are catalogued at pseudomonas.pseudomutant.com and pseudomonas.com [17]. Mutant strains were first grown in 96-well plates containing 150 μl of LB and incubated overnight at 37°C without shaking. Overnight cultures were transferred by 48/96 pin stamps into new 96-well plates containing 150 μl of sdH_2_O water, delivering ~10^7^ CFU of bacterial cells into each well. Gene expression (luminescence) in water was measured in black, 96-well, clear bottom microplates after inoculation into water (time 0) and at various time points using a Wallac Victor^3^ luminescence plate reader (PerkinElmer, USA). At each time point, plates were shaken for 5 s in a 0.10 mm-diameter orbital, followed by absorbance (600 nm) and luminescence (counts per second; CPS) readings in each of the wells. Gene expression was divided by the absorbance (CPS/OD_600_) to normalize for differences in cell density. Relative gene expression was determined by calculating the fold change in expression for each gene at each time point relative to the time zero (T0) time point. Strains harboring *lux* fusions in genes that demonstrated meaningful fold changes (>2 fold up or down regulated) were identified. Cluster analysis of gene expression was performed using TreeView and Cluster software.

## RESULTS

### *P. aeruginosa* is capable of long-term survival in water

PAO1 was grown overnight in LB medium, washed extensively, and inoculated into water at a concentration of 10^7^ CFU/ml and incubated at room temperature. Bacterial quantitation was performed to determine bacterial survival over time. The viability of PAO1 remained high over the course of 8 weeks (Fig. 1). Cell numbers (CFU/ml) on occasion declined modestly by 0.5 - 1 log from the initial inoculum, which might be due to the use of stationary phase cells as an inoculum, as a small percentage of stationary phase may be dead or dying. The initial decrease in viability may also be a result of lysis from osmotic shock (Fig. 1). Generally, viability remained at stable cell numbers for up to 8 weeks (Fig. 1), consistent with a previous study reporting *P. aeruginosa* survival in water for 145 days (20.7 weeks) [5].

**Figure 1.**
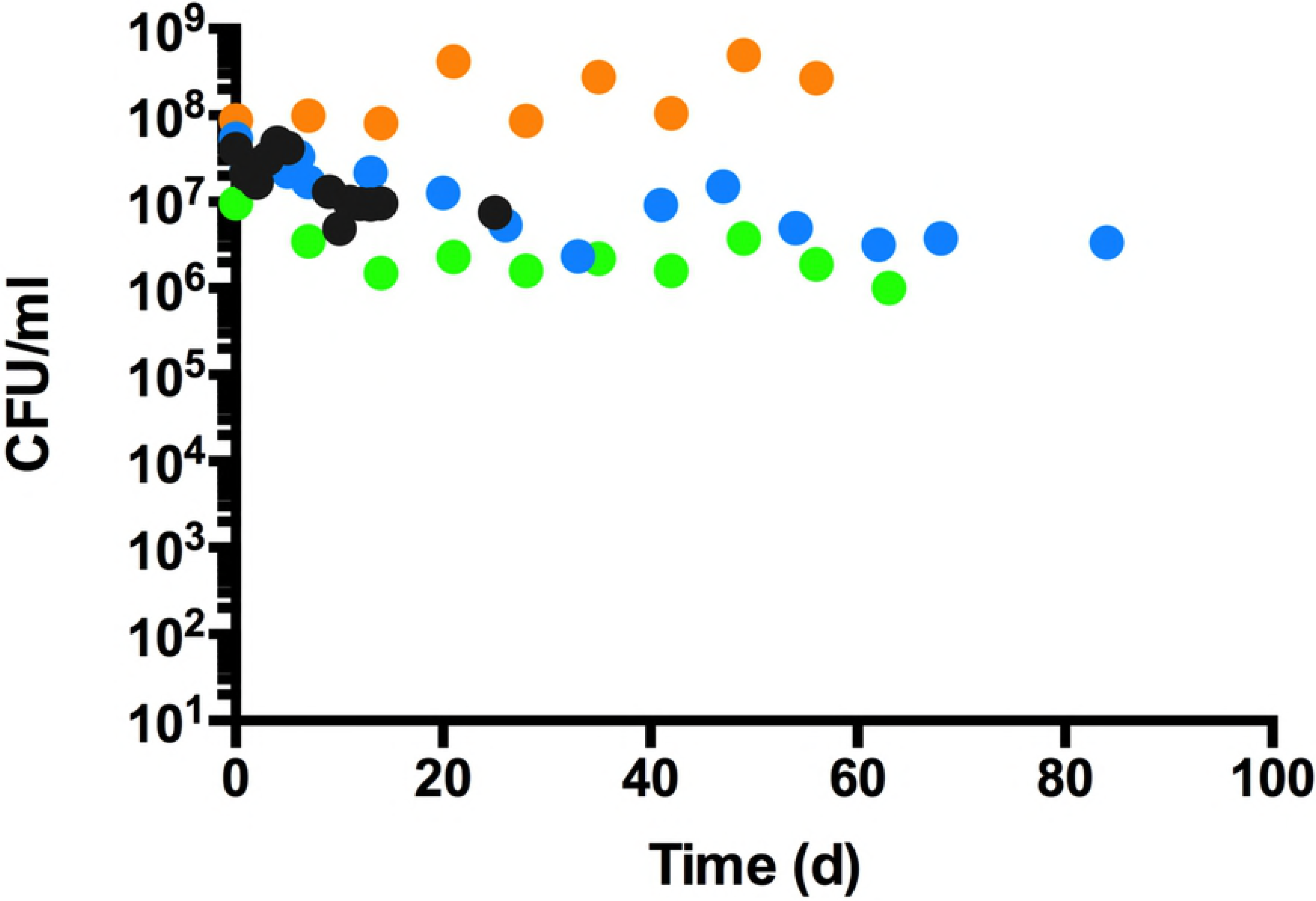
Survival of *P. aeruginosa* PAO1 incubated in water. Overnight cultures of PAO1 and mutants were washed thoroughly and inoculated into sterile water at a concentration of 10^7^ CFU/ml and incubated at room temperature. At each time point, aliquots were removed and plated on LB for direct bacterial counts. Each colour represents one of four trials and each value is the average of at least triplicate samples.

To increase the robustness of counting viable *P. aeruginosa* cells in water, we assessed the LIVE/DEAD staining patterns of cells in water using the DNA binding dyes SYTO 9 and propidium iodide (PI). While the majority of cells in water were a homogenous SYTO 9 population, there was also a subpopulation of PI stained cells (Fig. 2A). SYTO 9 is membrane permeable DNA stain that labels all cells green and PI is a membrane-impermeable, red DNA stain that was originally used as an indicator of non-viability. However, it is now recognized that PI is more likely an indicator of outer membrane disruption and increased permeability [18,19]. The numbers of total cells (green) and non-viable or membrane-compromised cells (red) were determined. In the early weeks, the proportion of PI-stained cells was low (6–20%), and increased to ~50% of cells at 4 weeks (Fig. 2C). The PI-stained proportion decreased in the later weeks to 15–30% of the population (Fig. 2C). In general, across all time points, ~75% of cells stained only with SYTO 9, and ~25% cells were staining with SYTO9 and PI, indicating this subpopulation was membrane-compromised and permeable to the DNA binding dye PI (Fig. 2D).

**Figure 2.**
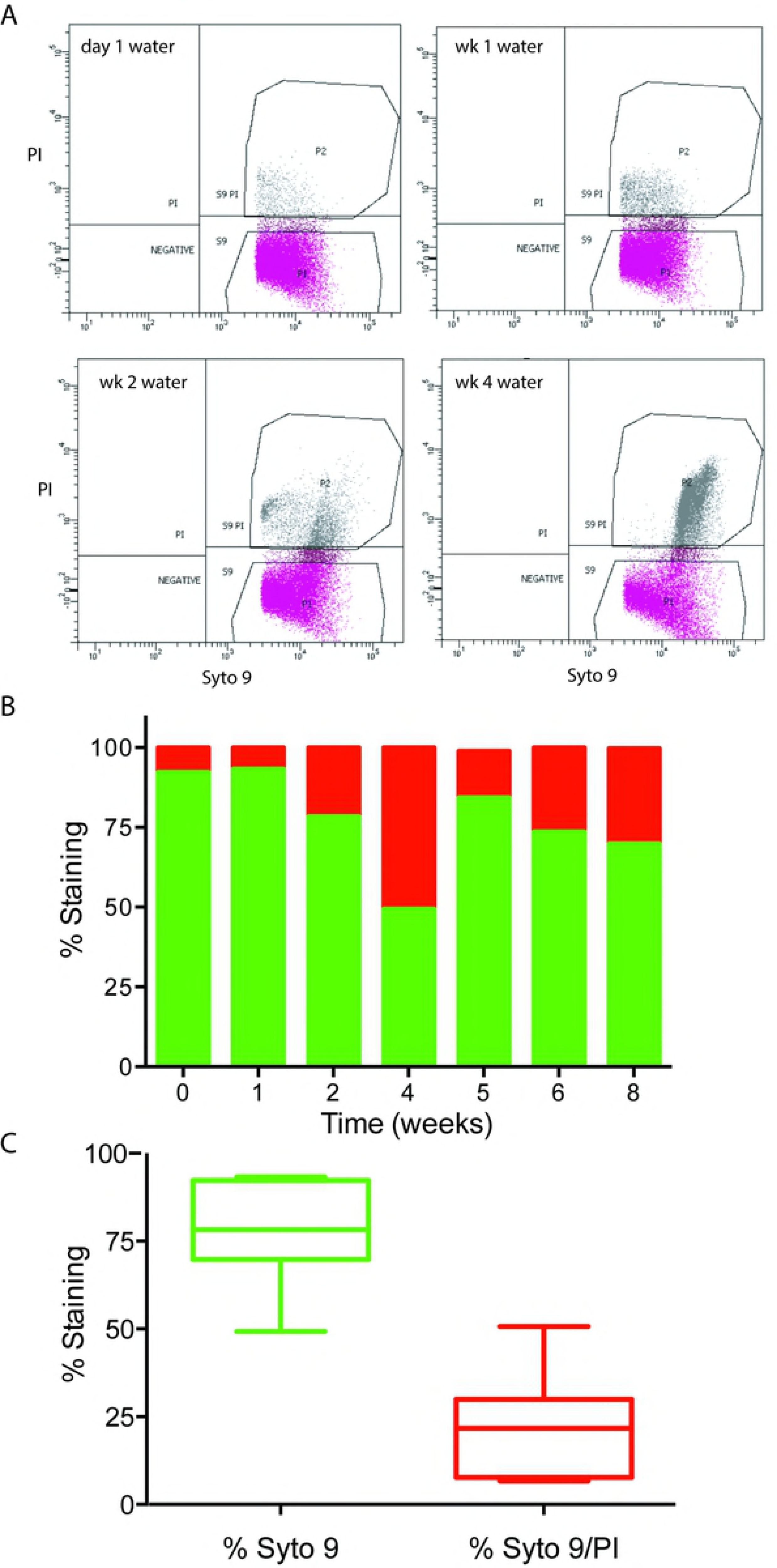
Scatter plots and bar graph representation of the populations of SYTO 9 and propidium iodide stained *P. aeruginosa* PAO1 in water. Strains were inoculated into sterile water at a concentration of 10^7^ CFU/ml and incubated at room temperature. A) Live and dead cell populations were subjected to LIVE/DEAD® staining every week and quantitated by flow cytometry. The quadrant labelled S9 refers to the cells that were stained with SYTO9 only, which are generally considered to be viable. The quadrant labelled S9PI refers to the cells that stained both with SYTO9 and PI, which are possibly dead or membrane-compromised, dormant cells. The time points are from day 1, week 1, week 2 and week 4 in water. Each panel represents a population of 50,000 cells per experiment. B) The percentage of SYTO 9 (green bars) and SYTO9/PI (red bars) stained cells is depicted over an 8-week time course. The values shown are the average S9 and S9PI counts recovered from triplicate flow cytometry samples. D) The box and whiskers plot demonstrates the overall proportion of SYTO 9 (green) only staining, compared to SYTO 9 and PI (red) staining populations of cells in water.

#### Fluorescence-activated cell sorting reveals that PI-stained cells are viable

To determine if PI-stained cells in water were viable, the SYTO 9 positive and SYTO 9/PI double positive population were separated and sorted into different tubes by FACS analysis. Next, the PI-stained populations were serially diluted and plated to determine if these cells were viable. The supposed ‘dead’ population of PI-stained cells indeed grew and was recovered after plating on LB agar, indicating that the PI-stained cells were membrane-damaged but viable. While monitoring the proportion of recoverable PI-staining cells, we observed that up to 100% of PI-stained cells were recovered after week 4 (Table 1). The percent viability was calculated by dividing the number of viable cells recovered on agar plates following serial dilution, by the number of PI-stained cells that were counted and sorted by FACS. At the early 1 week time point, it appears that over 50% of cells that stained with PI were non-viable, and it appears that *P.aeruginosa* undergoes a transition whereby the Pi-stained cells after 4 weeks are adapted to surviving in water and become fully viable (Table 1).

**Table 1.** The percentage of viable, propidium iodide-stained and overall ATP production of *P. aeruginosa* PAO1 in water.

| **Time (weeks)** | **% Viable cells<sup>a</sup>** | **ATP production (cps) <sup>b</sup>** |
| --- | --- | --- |
| 1 | 56.7 | 32161 |
| 4 | 100 | 1460 |
| 6 | 100 | 1334 |
| 8 | 100 | 1092 |

<sup>a</sup> Cells were FACS sorted that were dual positive for SYTO9 and PI, and plated for direct bacterial counts. <sup>b</sup> Cells (10<sup>7</sup> CFU in water) were incubated for 5 minutes at room temperature at a 1:1 ratio with BacTiter-Glo™ reagent (1 ml final volume) and luminescence as a measurement of ATP production was determined.

Having assessed viability of *P. aeruginosa* in water, we wanted to assess the overall metabolic activity. At various time points, a sample of PAO1 in water was taken and the ATP reagent from the BacTiter-Glo™ Microbial Cell Viability Assay from Promega was added at an equal volume. ATP production was measured in a luminometer as a function of counts per second (cps) of luminescence. ATP production decreased over time in the water samples from week 1 to 8, which demonstrates that the population reduced their overall metabolic activity (Table 1), yet were completely viable after 4 to 8 weeks (Fig. 1, Table 1).

#### *P. aeruginosa* DNA is intact and may be highly condensed in water

Further flow cytometry experiments were performed using PicoGreen®, another viability stain that measures vitality based on intact, dsDNA content. For this experiment, mid-log phase cultures of PAO1 were inoculated into water for incubation, aliquots were removed over time, stained with PicoGreen® and subjected to flow cytometry. Overall, PAO1 was confirmed to be viable in water as the cells stained with PicoGreen®, resulting in strong green fluorescence. However, these experiments also revealed that PAO1 fluoresced more in water compared to LB, and became more fluorescent over time, with a 100-fold increase in peak fluorescence comparing cells in LB (10^3^) to cells inoculated into water (10^5^) (Fig. 4). By 3 and 4 weeks in water, the peak fluorescence was found to be >10^5^. The PicoGreen® fluorescence profile was generally lower and more dispersed when stationary phase cells were incubated in water, as this population contains both living and dying cells, and likely a mixture of dsDNA and degraded DNA (data not shown). During the 4 week incubation period in water, the picogreen fluorescence profile transitioned to a homogenous, high fluorescence profile, possibly reflecting a transition towards a stable population of dormant cells and more intact dsDNA. The high fluorescence profile of cells in water for 4 weeks may also suggest that the DNA structure became more tightly supercoiled and therefore more highly condensed (Fig. 4). In addition, the side-scatter values (SSC) decreased over time in water compared to LB (Fig. 4), which may indicate the transition to a smaller cell size, which was observed in microscopy (Fig. 3).

**Figure 3.**
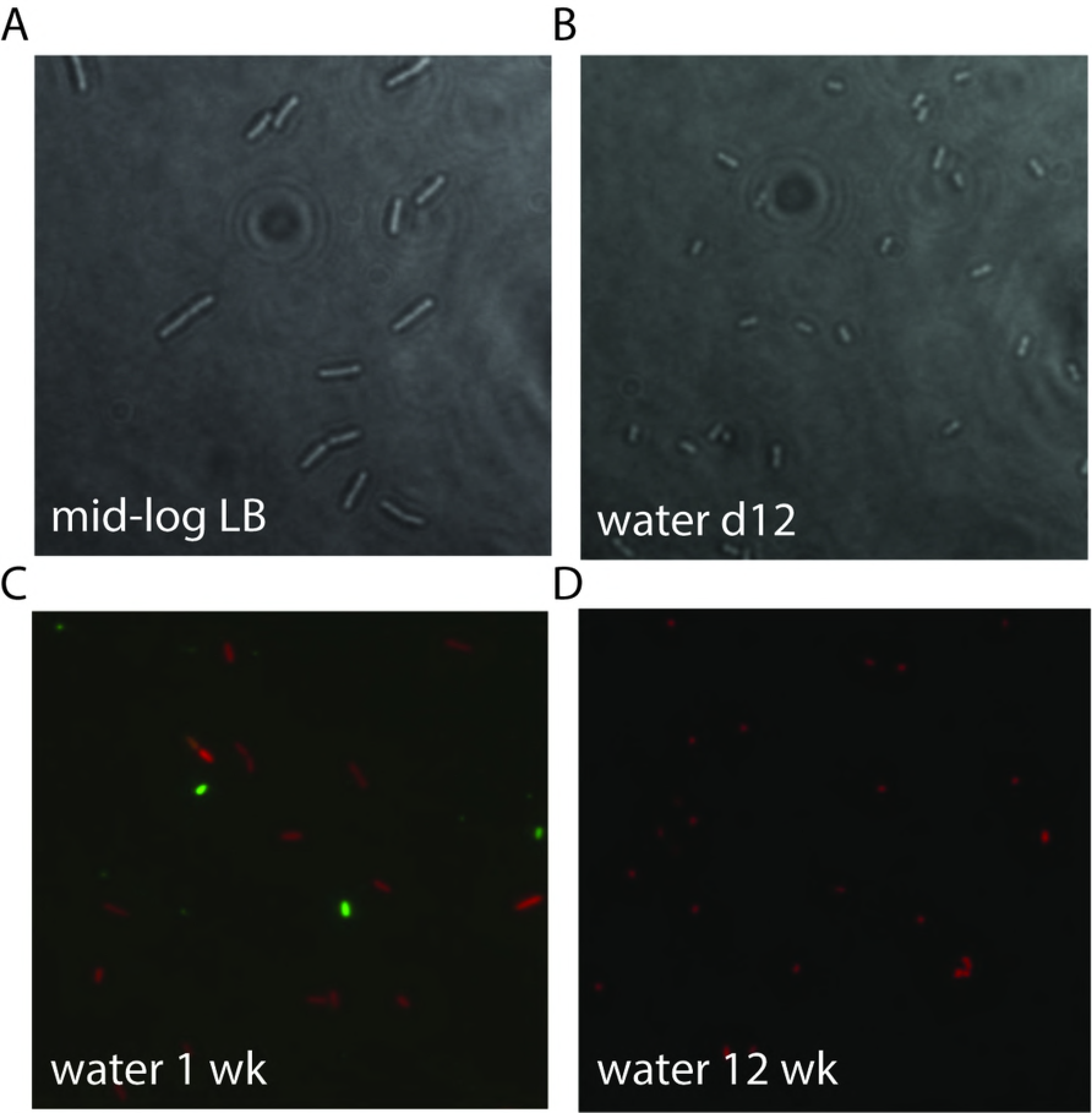
Phase-contrast and fluorescence microscopy of *P. aeruginosa* PAO1 in water. A) Mid-log cells of *P. aeruginosa* under phase contrast. B) *P. aeruginosa* incubated in water for 12 days under phase contrast. C) LIVE/DEAD® staining results of *P. aeruginosa* PAO1 following incubation in water for 1 and D) 12 weeks. Cells were grown to mid-log in LB, washed, and added to sdH_2_O at a concentration of 10^7^ CFU/ml and incubated at room temperature. Cells were added to agarose beds on glass slides and visualized on a Leica microscope.

**Figure 4.**
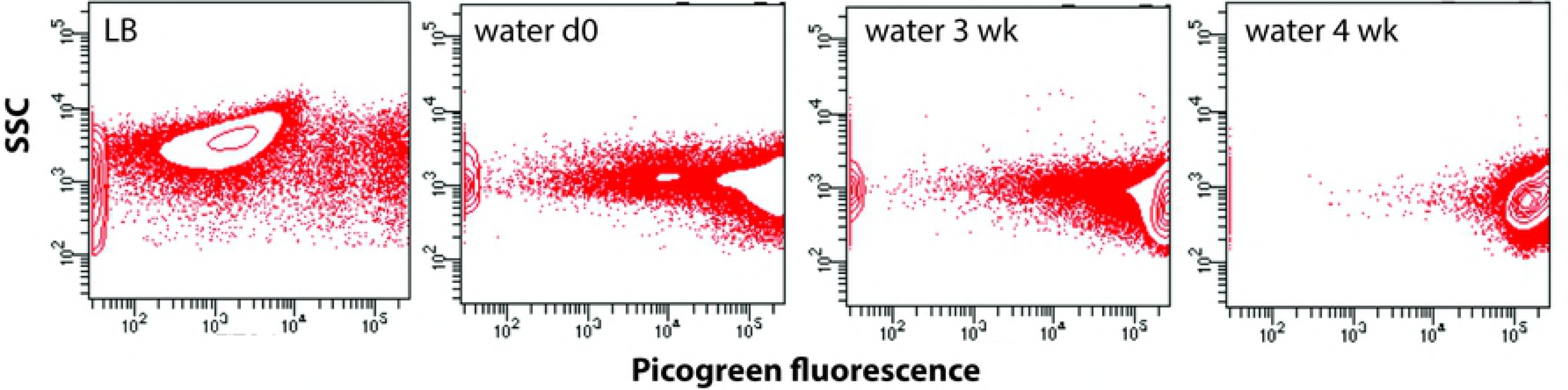
Pico Green staining and flow cytometry analysis of *P. aeruginosa* PAO1 in water. Log phase cultures of PAO1 were inoculated into water for incubation. Each sample was stained with Quant-iT™ PicoGreen® and subjected to flow cytometry to analyze the dsDNA content in cells during long-term incubation in water. Cells from mid-log PAO1 cultures were compared to cells in water at day 0, 3 weeks and 4 weeks. Each panel represents a population of 50,000 cells per experiment.

#### The PAO1 outer membrane is tolerant to polymyxin B after prolonged incubation in water

Outer membrane permeability of PAO1 was assessed using a previously established method [13]. Cells were pretreated with sodium azide, which blocks active efflux mechanisms, and then exposed to 1-N-phenylnaphthylamine (NPN). The baseline NPN fluorescence in sodium azide-treated cells reflects the amount of NPN that penetrates the hydrophobic regions of the membrane, where NPN becomes fluorescent. Prior to the addition of polymyxin B to disrupt the outer membrane, the baseline of membrane permeability decreases over time during incubation in water, with very low NPN fluorescence in cells after 3 weeks in water (Fig. 5A).

**Figure 5.**
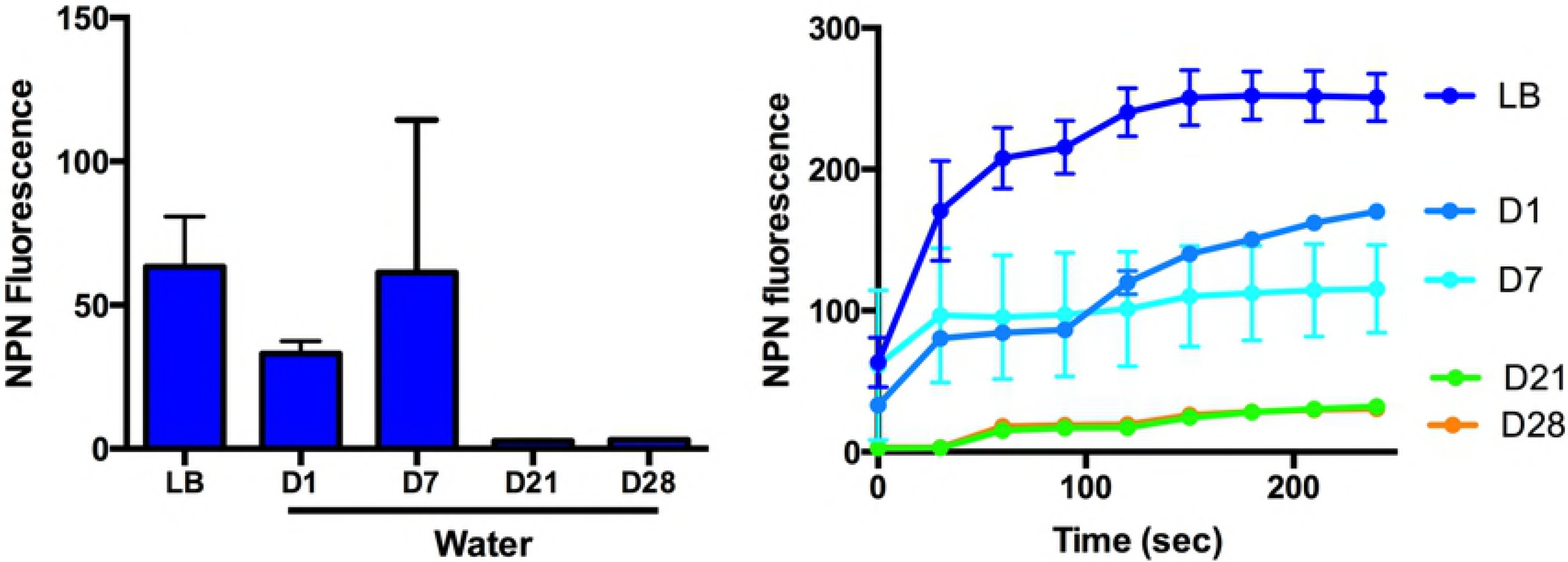
Outer membrane permeability and polymyxin B tolerance of *P. aeruginosa* PAO1 incubated in water. A) The baseline of outer membrane permeability was measured as a function of 1-N-phenylnaphthylamine (NPN) uptake and subsequent fluorescence in relative light units (RLU). Log phase cultures of PAO1 were prepared and inoculated into water for incubation. After 1, 7, 21, and 28 days incubation, cells were treated with sodium azide, an active efflux inhibitor. B) After NPN addition, polymyxin B was added to disrupt the outer membrane and increase NPN uptake into the hydrophobic environment of the envelope. The tolerance to polymyxin B treatment was compared between mid-log LB cultures and cells incubated in water for up to 4 weeks. Values shown are the average and standard error of triplicate samples.

After the addition of polymyxin B, the outer membrane of mid-log phase PAO1 grown in LB was disrupted leading to a rapid increase in NPN incorporation and fluorescence (Fig. 5B). The response of PAO1 after 1 day in water was similar but incorporated less NPN after polymyxin B treatment, and the even less membrane damage at day 7. Cells incubated in water for 3 and 4 weeks demonstrated a polymyxin B resistance phenotype and showed very little NPN fluorescence (Fig. 5). This data suggests that the outer membrane of PAO1 is altered in this low nutrient environment, resulting in a reduction in membrane permeability and an increase in tolerance to disruption by the antimicrobial peptide polymyxin B.

#### Differential expression patterns of *P. aeruginosa* PAO1 genes in water

To assess the gene expression patterns of PAO1 in water, we incubated a previously described collection of mini-Tn5-*luxCDABE* transposon mutants of *P. aeruginosa* [16]. This transposon mutant library of *P. aeruginosa* PAO1 is a collection of random transposon mutants, where each mini-Tn*5-lux* insertion creates an active transcriptional *luxCDABE* fusion if inserted into the same orientation as the gene. The mini-Tn5-*luxCDABE* library in PAO1 contains about ~2,500 mutants with mapped Tn insertion sites, and ~1,400 of these were transcriptional *lux* fusions [16]. This entire collection of ~2500 mini-Tn*5-lux* mutants was screened for mutants with impaired survival in water, however no mutants were identified that had significant defects in survival during longterm incubation in water. This observation suggests that single gene knockouts cannot result in survival defects, or that the genes of interest were not present in our mutant library.

The collection of ~1,400 mapped transcriptional *lux* fusions were inoculated into 96-well plates containing sterile water (10^7^ CFU/well) and incubated at room temperature. The gene expression patterns were determined by measuring absorbance (OD_600_) and luminescence (CPS) at 0, 0.3, 1, 3, 5, 7, 13, 20, 26, and 34 days. The data was analyzed by dividing luminescence by absorbance to correct for differences in cell density, and the fold changes of expression were determined by comparing all values to time 0 (Sup Table 1). Cluster analysis of gene expression was performed to assess the overall trends in gene expression.

The vast majority of genes were repressed (orange) during incubation in water. However, there were sub-groups of genes that were induced (blue) during incubation in water (Fig. 7, Sup Fig. 1-4). Four clusters of induced genes were identified: Cluster A, containing genes that were found to be induced very late, around one month in water; Cluster B, genes that were induced throughout the time course; Cluster C, genes that were induced at early time points and then repressed in later time points; and Cluster D, genes that were expressed late (close to one month in water) (Sup. Fig. 1-4).

**Figure 6.**
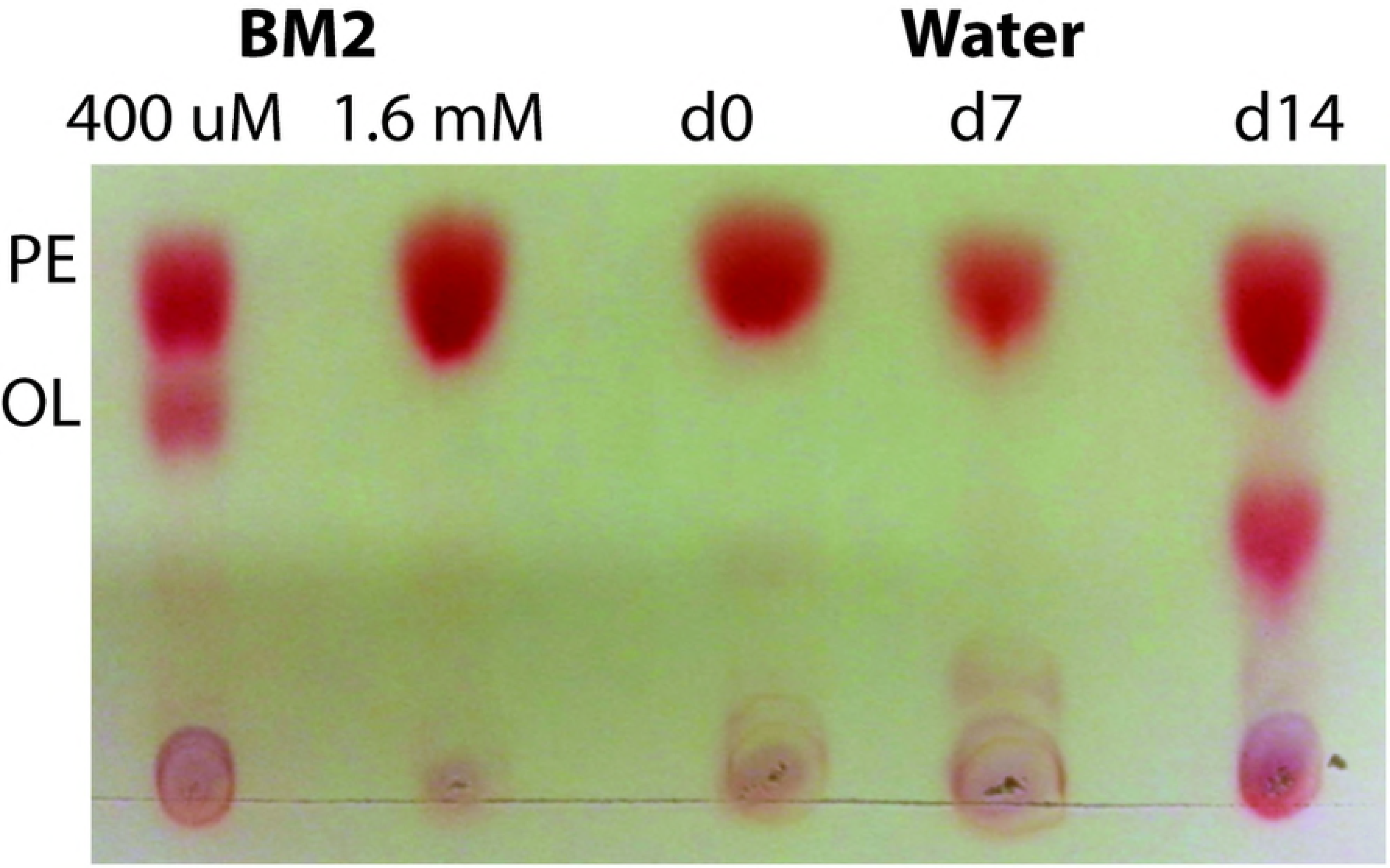
Ninhydrin detection of amino group containing lipids of PAO1 cells following incubation in water. Total lipids were extracted, separated by thin layer chromatography, and sprayed with ninhydrin to visualize the amino group-containing lipids. Lipid samples of water cultures at day 0, day 7 and day 14 were run on TLC plate, alongside controls of lipids extracted from cultures grown in BM2-defined media with limiting (400 μM) and high phosphate (1.6 mM) conditions. The positions of the primary membrane lipid phosphatidylethanolamine (PE) and the unique ornithine lipid (OL) species that is produced under phosphate limitation [15] are indicated.

**Figure 7.**
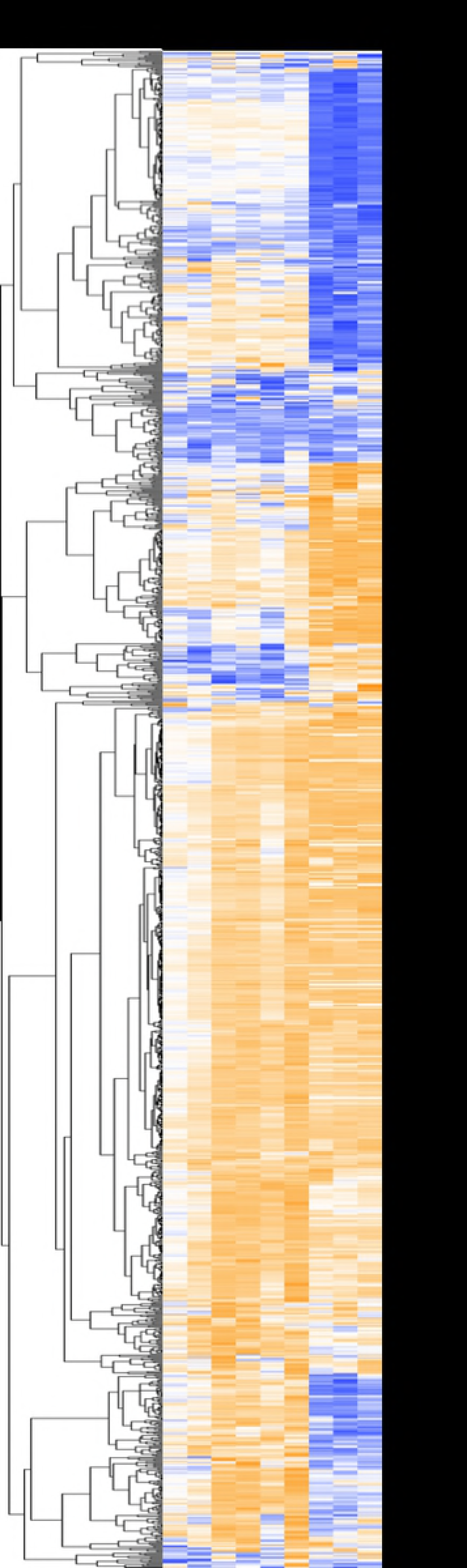
Cluster analysis of gene expression of *P. aeruginosa* genes in response to water. The PAO1 mini-Tn5-*luxCDABE* mutant library containing 1369 transcriptional *lux* fusion strains was inoculated into water in black 96 well microplates and incubated at room temperature. At each time point the optical density (OD_600_) and luminescence (counts per second) was measured. Gene expression (CPS) readings were taken at day 0, 0.3, 1, 3, 5, 7, 13, 20, 26, and 34. Luminescence was divided by absorbance and fold changes were calculated based on the change in expression (CPS/OD_600_) compared to time 0. Cluster analysis was performed using Tree View and Cluster 3.0 software. Orange indicates repression, and blue indicates induced expression, relative to the time zero point. Genes with no change in expression are in white. Black bars highlight clusters of genes that are induced late (A, D), throughout (B) or early (C) in the 34 day time period of incubation in water.

Some of the repressed genes were identified as those involved in DNA replication. Genes such as *holB* and *sss*, encoding for DNA polymerase III and a site-specific recombinase, were induced early on, but then repressed for the remainder of the experiment, indicating that DNA replication slows over time in water. Other DNA replication genes such as *recQ* and *ruvA*, were repressed in water as well. However, the *sbc* genes (*sbcB* and *sbcD*), which encode for exonucleases (DNases) were induced in water. Genes encoding for proteins responsible for mismatch repair (*micA*) and those involved in nucleotide excision repair (uvr) were induced. In addition, *polA*, the gene encoding for DNA polymerase I, which has an exonuclease activity (3’–5’ and 5’–3’) required for excision repair was induced in water as were DNA binding proteins PA3940 and PA4704 (*cbpA*), and integration host factor *himA*. These results suggest that DNA repair and nucleoid condensation may occur to protect the DNA in unfavorable conditions. Induction of nucleoid-associated proteins (NAPs) such as *cbpA* and *himA* correlates with the increased fluorescence observed in PicoGreen® staining.

Many metabolic genes were repressed in water, which is consistent with the reduction in ATP production over time (Table 1), but some genes were induced that indicate a shift to utilizing alternate sources of energy to persist. Genes required for fatty-acid oxidation and involved in fatty-acid and phospholipid metabolism (*foaB*, *fadH1*, *fadH2*) were induced very late and may explain the utilization of the phospholipid membrane for energy and the reduction in cell size after one month in water. Many amino acid uptake (PA4911, PA4072, *oprD*) and catabolism genes (*pepA*, *aruB*, *phhC*, *amaB*, *gcdH*, *ilvD*) were also induced, pointing to increased uptake and degradation of amino acids as a source of nutrients, possibly from the subset of dying cells in the population. Other expressed genes (*exaA*, *zdhB*), responsible for the utilization of alternative energy sources (alcohols, xanthine, purines, pyrimidines, pterins, and aldehyde substrates) were up-regulated as well.

Genes that were consistently expressed or induced over two months in water were those that may be involved in maintaining the electrochemical gradient, or proton-motive force (PMF) of the membrane (*cycH*, *cyoB*, *cyoC*, and *ccpR*), a number of transport and efflux-related genes (*oprD*, *spuF*, *mexD*, PA0397, PA0450, PA4126, PA3840), many transcriptional regulators (PA0163, PA3782, PA5179, PA0272, *dnr*), and genes encoding for sensor/response regulators (*retS*, *phoB*, *pilS*, PA4293). The expression patterns of these genes demonstrates a requirement for PAO1 in nutrient depleted conditions to maintain the PMF in order for the organism to synthesize minimal amounts of ATP by using alternative energy sources, to preserve essential cell components, transport substrates, and respond to the environment, all through coordinated transcriptional control of cellular processes. The presence of a constant PMF was also observed in flow cytometry experiments using Redox Sensor Green, a cell viability stain based on the presence of membrane potential, as PAO1 cells were successfully stained after 6 days in water (data not shown).

There were a number of genes of interest that had notable gene expression patterns in water over time, suggesting a specific role in persistence. Genes associated with adaptation and protection such as *inaA*, and *cyaA* were both induced early on in water and then repressed at later time points. Flagella genes (*flgJ*, *fliM*, *flgK*, *flhA*, *fliC*) were induced at later time points, as were the type VI secretion genes (*tsel*, *vgrG*).

Supplemental Table 1 lists the identity of each gene, the raw expression (CPS) and fold change in gene expression at every time point relative to time zero. The global pattern of gene repression suggests that many cellular processes are turned off during prolonged incubation in water, however some genes were found to be induced at particular time points or at much later time points, possibly indicating a specific role in water survival and a highly coordinated response.

#### *P. aeruginosa* alters the phospholipid composition of its membrane when dormant

The longer *P. aeruginosa* remained in water, viability increased, and the outer membrane became more impermeable to the hydrophobic dye NPN and more tolerant to polymyxin B. To further test the hypothesis that the membrane undergoes changes during incubation in water, we analyzed the total phospholipid content of cells in water. Thin layer chromatography (TLC) was used to separate the total lipid extracts from PAO1, following incubation in water. Cells were incubated in water as described above and samples were taken at day 0, day 7, and day 14 and run on a TLC plate with lipids extracted from control cells grown in low and high phosphate concentrations. In the presence of limiting phosphate, *P. aeruginosa* produces a unique ornithine lipid that lacks phosphate in the head group, as a mechanism of adapting to limiting phosphate [15], as a substitute for the primary lipid in the PAO1 envelope, phosphatidylethanolamine. PAO1 produced a novel lipid species following incubation in water for 14 days, which was not ornithine lipid, indicating that *P. aeruginosa* produces a different membrane phospholipid content during incubation in water (Fig. 6).

## DISCUSSION

*Pseudomonas aeruginosa* is capable of long-term survival without nutrients by existing in a dormant state. Despite staining with propidium iodide, cells sorted by FACS analysis were plated on LB agar and found to 100% viable within 4 weeks (Fig. 2, Table 1). The cell-impermeable DNA stain propidium iodide (PI) was originally thought of as a stain for dead cells, but in agreement with other findings [18], we demonstrate here that PI-stained cells can be sorted and recovered as viable growing colonies. PI is therefore better described as an indicator of membrane damage, rather than bacterial death. During long-term survival in water, *P. aeruginosa* displayed several adaptations that are consistent with dormancy. Cells in water had decreased metabolic activity, as determined by measuring ATP production and by a general repression trend in the gene expression patterns of a large number of transcriptional *lux* fusions. Cell shape converted from a rod to a coccoid shape, the phospholipid content changed, and the outer membrane demonstrated a decreased permeability and increased tolerance to polymyxin B disruption.

Although the majority of PAO1 genes were suppressed in water, a number of genes were found to maintain or have induced expression at some point in the time course of long-term survival in water. Since we were unable to recover single transposon mutants that died during longterm incubation in water, it appears that the adaptation to surviving in water is complex and involves more than a single gene. Given the substantial proportion of induced genes, there does appear to be an active and complex process of differentiation. It may be that multiple genes contribute to survival in water and therefore it is unlikely to identify single mutants with survival defects. The mutant library used here is not a saturating collection of mutants, and the mutants of interest may not be present in this library [16,20]. Future experiments will employ other genome-wide methods to attempt to identify a specific mechanism and genes required for long-term survival in water.

The gene expression profile of *P. aeruginosa* in water validates a coordinated response by the organism in the transition to dormancy. Most of the genes were repressed over time indicating a reduction in many cellular and metabolic processes, but a number of specific genes were induced throughout, or at certain time points, suggesting an importance for these genes in survival and maintenance of a dormant state. Amino acid, fatty acid, and phospholipid metabolism genes were induced and these compounds likely become alternative energy sources in the absence of nutrients [21]. It has been shown that growth-arrested bacteria utilize membrane phospholipids as an alternative energy source, which leads to a reduction in cell size and volume and promotes transport of substrates [9]. This has been noted in both *E. coli* and *Vibrio cholerae* [22,23]. The cell size of *P. aeruginosa* significantly decreased over time in water (Fig. 3), and we also observed an increase in expression of a number of transport genes (Sup Table 1).

Genes in the library required for DNA replication were repressed over time, but those involved in DNA repair and DNA packaging were induced. It is likely that DNA repair and packaging are needed to ensure that DNA is protected and the fidelity of DNA is maintained until the cell is in more favorable conditions for replication. The induction of genes encoding for nucleoid-associated proteins (NAPs) such as CpbA and HimA, involved in nucleoid condensation in other species, was notable as this correlates with the observed increase in fluorescence of PAO1 in water over time stained with the dsDNA dye PicoGreen® (Fig. 4). *E. coli* in stationary phase has been shown to possess a condensed nucleoid due to NAPs, which is thought to protect against DNA damage and confer a survival advantage [9,24].

Gene expression results also indicated that genes involved in maintenance of the proton-motive force (PMF), efflux pumps, sensor-response regulators, and other transcriptional regulators were required for *P. aeruginosa* in water. Maintenance of the PMF is very likely necessary to allow the organism to make ATP using alternative energy sources, transport substrates, support motility, and respond to the environment [9]. It has been shown that bacteria in growth arrest need to preserve the PMF to enable these functions in addition to maintaining the essential macromolecular components of the cell [25,26]. Many flagellin genes were also induced in water, indicating that flagella are needed for biofilm formation, and possibly for chemotaxis to a nutrient source [27,28]. Large aggregates, likely biofilms, were observed by microscopy and detected in flow cytometry in PAO1 after 2 months in water (data not shown).

Genes involved in adaptation and protection were induced early on in water. The role of these genes will be investigated in future studies. Type VI secretion genes were significantly induced after one month in water and they may be needed for a competitive or protective advantage, as well as biofilm formation (*tse1*), but further studies will be required to determine the role of the type VI secretion system in dormancy [29,30]. The differential expression of PAO1 genes in water using the mini-Tn*5-lux* library provides significant insight into the complexity of the response of the organism to this environment and points to a number of potential mechanisms required for survival in water.

Non-sporulating bacteria undergo a reversible state of low metabolic activity without replication to persist in unfavorable environmental conditions [31,11]. Previous studies have indicated that other non-spore forming bacteria such as *E. coli* and *Klebsiella pneumoniae* are capable of dormancy under environmental stress and that this is a reversible phenomenon [32]. Dormancy in *Mycobacterium tuberculosis* has been well documented [33]. Dormant cells have also been referred to as persister cells because they are able to resist the effects of antibiotics [34]. Although traditionally persister cells and dormancy have been considered to be separate phenomena, because persister cells result after exposure to high doses of antibiotics, studies have shown that dormancy may be the best model for persister cells [35,36]. Aside from exposure to antibiotics, other inducers of persistence or dormancy in bacteria may be stress or starvation responses [37–40]. In general, some mediators of persistence in bacteria have been shown to be the SOS response genes, TisB toxin, the RelA protein, and the HipB toxin [37,48]. In addition, high persister cells (hip) are often found within a biofilm [37]. Bacterial persistence is a major issue when dealing with infectious diseases as these cells are resistant to antibiotics [34]. The phenomenon of dormancy and persistence has been investigated in *P. aeurginosa*, primarily in terms of antibiotic resistance, biofilm formation, and resistance to chemicals [37,42,43] and some novel persister genes have been identified [44].

Similar to our results, *Vibrio cholerae* has been shown to shift to a persister phenotype in water [45]. When *V. cholerae* was introduced into filter-sterilized lake water the cells displayed characteristics of persister cells and were culturable for >700 days. Interestingly, these authors also observed that the cells became smaller and formed aggregates over time in water, similar to what was observed in this study. The authors concluded that nutrient stress can induce a persister phenotype in *V. cholerae* in environmental reservoirs, which results in epidemics of the disease when nutrients such as phosphate become more available in the environment. We are interested in determining the nutrient threshold required to revert persister cells in *P. aeruginosa* to vegetative cells.

*P. aeruginosa* can also be considered a model organism for the study of diverse bacterial mechanisms that contribute to bacterial persistence. The ability of *P. aeruginosa* to survive long term in water, and to be recovered from drains, sinks, and water pipes, makes it a reservoir for infectious disease. Since this organism is readily transferred into the hospital where it causes infection, it is important to understand how this organism survives. This will contribute to solutions for the prevention of infection. Finally, determining the mechanism for survival in water will be beneficial for understanding how other microorganisms may persist in similar conditions.

## ACKNOWLEDGEMENTS

This work was funded by the Athabasca University Academic Research Fund. Shawn Lewenza held the Westaim-ASRA Chair in Biofilm Research. We would like to thank Dr. Richard Moore for insights into water survival experiments and assisting with experimentation.

## REFERENCES

1. Jorgensen F, Bally M, Chapon-Herve V, Michel G, Lazdunski A, Williams P, Stewart GS. 1999. RpoS-dependant stress tolerance in Pseudomonas aeruginosa. Microbiol. 1999;145;835–844. pmid:10220163

2. Wolfgang MC, Kulasekara BR, Liang X, Boyd D, Wu K, Yang Q, et al. Conservation of genome content and virulence determinants among clinical and environmental isolates of Pseudomonas aeruginosa. PNAS. 2003;100(14);8484–8489. pmid:12815109

3. Driscoll JA, Brody SL, Kollef MH. The epidemiology, pathogenesis and treatment of Pseudomonas aeruginosa infections. Drugs. 2007;67(3);351–368. pmid:17335295

4. Kramer A, Schwebke I, Kampf G. How long do nosocomial pathogens persist on inanimate surfaces? A systematic review. BMC Infectious Dis. 2006;6;130. pmid:16914034

5. Moore RA, Tuanyok A, Woods DE. Survival of Burkholderia pseudomallei in water. BMC Res Notes. 2008;1;11.

6. Hemme CL, Tu Q, Shi Z, Qin Y, Gao W, Deng Y, et al. Comparative metagenomics reveals impact of contaminants on groundwater microbiomes. Front Microbiol. 2015;6;1205. pmid:26583008

7. Trautmann M, Lepper PM, Haller M. Ecology of Pseudomonas aeruginosa in the intensive care unit and the evolving role of water outlets as a reservoir of the organism. Am J Infect Control 2005;33(5Suppl.1);S41–49. pmid:15940115

8. Aumeran C, Paillard C, Robin F, Kanold J, Baud O, Bonnet R. Pseudomonas aeruginosa and Pseudomonas putida outbreak associated with contaminated water outlets in an oncohaematology paediatric unit. J Hosp Infect. 2007;65(1);47–53. pmid:17141370

9. Bergkessel M, Basta DW, Newman DK. The physiology of growth arrest: uniting molecular and environmental microbiology. Nat Rev Microbiol. 2016;14(9);549–562.

10. Lewis K. Persister cells, dormancy and infectious disease. Nat Rev Microbiol. 2007;5(1);48–56. pmid:17143318

11. Lennon JT, Jones SE. Microbial seed banks: the ecological and evolutionary implications of dormancy. Nat Rev Microbiol. 2011;9(2);119–130. pmid:21233850

12. Gefen O, Fridman O, Ronin I, Balaban NQ. Direct observation of single stationary-phase bacteria reveals a surprisingly long period of constant protein production activity. PNAS. 2014; 111(1);556–561. pmid:24344288

13. Lo B, Grant C, Hancock RE. Use of the fluorescent probe 1-N-phenylnaphthylamine to study the interactions of aminoglycoside antibiotics with the outer membrane of Pseudomonas aeruginosa. Antimicrob. Agents Chemother. 1984;26(4);546–551. pmid:6440475

14. Johnson L, Mulcahy H, Kanevets U, Shi Y, Lewenza S. Surface-localized spermidine protects the Pseudomonas aeruginosa outer membrane from antibiotic treatment and oxidative stress. J Bacteriol. 2012;194(4);813–826. pmid:22155771

15. Lewenza S, Falsafi R, Bains M, Rohs P, Stupak J, Sprott GD, et al. The olsA gene mediates the synthesis of an ornithine lipid in Pseudomonas aeruginosa during growth under phosphate-limiting conditions, but is not involved in antimicrobial peptide susceptibility. FEMS Microbiol Lett. 2011;320(2);95–102. pmid:21535098

16. Lewenza, S, Falsafi RK, Winsor G, Gooderham WJ, McPhee JB, Brinkman FS, et al. Construction of a mini-Tn5-luxCDABE mutant library in Pseudomonas aeruginosa PAO1: a tool for identifying differentially regulated genes. Genome Res. 2005;15;583–589. pmid:15805499

17. Winsor GL, Griffiths EJ, Lo R, Dhillon BK, Shay JA, Brinkman FS. Enhanced annotations and features for comparing thousands of Pseudomonas genomes in the Pseudomonas genome database. Nucleic Acids Res. 2016;4;44(D1);D646–653. pmid:26578582

18. Shi L, Günther S, Hübschmann T, Wick LY, Harms H, Müller S. Limits of propidium iodide as a cell viability indicator for environmental bacteria. Cytometry A. 2007;71(8);592–598. pmid:17421025

19. Halverson TW, Wilton M, Poon KK, Petri B, Lewenza S. DNA is an antimicrobial component of neutrophil extracellular traps. PLoS Pathog. 2015;11(1); e1004593. pmid:25590621

20. Stover CK, Pham XQ, Erwin AL, Mizoguchi AD, Warrener P, Hickey MJ, et al. Complete genome sequence of Pseudomonas aeruginosa PAO1, an opportunistic pathogen. Nature. 2000;406(6799);959–964. pmid:10984043

21. Daniel J, Deb C, Dubey VS, Sirakova TD, Abomoelak B, Morbidoni HR, et al. Induction of a novel class of diacylglycerol acyltransferases and triacylglycerol accumulation in Mycobacterium tuberculosis as it goes into a dormancy-like state in culture. J Bacteriol. 2004;186(15);5017–5030. pmid:15262939

22. Farewell A, Diez AA, DiRusso CD, Nystrom T. Role of the Escherichia coli FadR regulator in stasis survival and growth phase-dependent expression of the uspA. fad, and fab genes. J Bacteriol. 1996;178(22);6443–6450. pmid:8932299

23. Hood MA, Guckert JB, White DC, Deck F. Effect of nutrient deprivation on lipid, carbohydrate, DNA, RNA, and protein levels in Vibrio cholerae. Appl Environ Microbiol. 1986;52;788–793. pmid:2430523

24. Wolf SG, Frenkiel D, Arad T, Finkel SE, Kolter R, Minsky A. DNA protection by stress-induced biocrystallization. Nature. 1999;400(6739);83–85. pmid:10403254

25. Koch AL. Microbial physiology and ecology of slow growth. Microbiol Mol Biol Rev. 1997;61;305–318. pmid:9293184

26. Nystrom T, Gustavsson N. Maintenance energy requirement: what is required for stasis survival of Escherichia coli? Biochim Biophys Acta. 1998;1365;225–231.

27. Geesy GG, Morita RY. Capture of arginine at low concentrations by a marine psychrophilic bacterium. Appl Environ Microbiol. 1979;38;1092–1097. pmid:16345475

28. Klausen M, Heydorn A, Ragas P, Lambertsen L, Aaes-Jorgensen A, Molin S et al. Biofilm formation by Pseudomonas aeruginosa wild type, flagella, and type IV pili mutants. Mol Microbiol. 2003;48(6);1511–1524. pmid:12791135

29. Mougous JD, Cuff ME, Raunser S, Shen A, Zhou M, Gifford CA, et al. A virulence locus of Pseudomonas aeruginosa encodes a protein secretion apparatus. Science. 2006;312(5779);1526–1530. pmid:16763151

30. Southey-Pillig CJ, Davies DG, Sauer K. Characterization of temporal protein production in Pseudomonas aeruginosa biofilms. J Bacteriol. 2005;187(23);8114–8126. pmid:16291684

31. Kaprelyants AS, Gottschal JC, Kell, DB. Dormancy in non-sporulating bacteria. FEMS Microbiol Rev. 1993;10(3-4);271–285. pmid:8318260

32. Sachidanandham R, Yew-Hoong Gin K. A dormancy state in nonspore-forming bacteria. Appl Microbiol Biotechnol. 2009;81;927–941. pmid:18815783

33. Boon C, Dick T. How Mycobacterium tuberculosis goes to sleep: the dormancy survival regulator DosR a decade later. Future Microbiol. 2012;7(4);513–518. pmid:22439727

34. Rotem E, Loinger A, Ronin I, Levin-Reisman I, Gabay C, Shoresh N, et al. Regulation of phenotypic variability by a threshold-based mechanism underlies bacterial persistence. PNAS. 2010;107(28);12541–12546. pmid:20616060

35. Wood TK, Knabel SJ, Kwan BW. Bacterial persister cell formation and dormancy. Appl Environ Microbiol. 2013;79(23);7116–7121. pmid:24038684

36. Kell D, Potgieter M, Pretorius E. Individuality, phenotypic differentiation, dormancy and ‘persistence’ in culturable bacterial systems: commonalities shared by environmental, laboratory, and clinical microbiology. F1000 Res. 2015;4;179. pmid:26629334

37. Lewis K. Persister cells. Annu Rev Microbiol. 2010;64;357–372. pmid:20528688

38. Fung DKC, Chan EWC, Chin ML, Chan RCY. Delineation of a bacterial starvation stress response network which can mediate antibiotic tolerance development. Antimicrob Agents Chemother. 2010;54(3);1082–1093. pmid:20086164

39. Murakami K, Ono T, Viducic D, Kayama S, Mori M, Hirota K, et al. Role for the rpoS gene of Pseudomonas aeruginosa in antibiotic tolerance. FEMS Microbiol Lett. 2005;242;161–167. pmid:15621433

40. Viducic D, Ono T, Murakami K, Susilowati H, Kayama S, Hirota K, et al. Functional analysis of spot, relA, and dksA genes on quinolone tolerance in Pseudomonas aeruginosa under nongrowing condition. Microbiol Immunol. 2006;50;349–357. pmid:16625057

41. Lewis K. Multidrug tolerance of biofilms and persister cells. Curr Top Microbiol Immunol. 2008;322;107–131. pmid:18453274

42. Harrison JJ, Turner RJ, Ceri H. Persister cells, the biofilm matrix and tolerance to metal cations in biofilm and planktonic Pseudomonas aeruginosa. Environ Microbiol. 2005;7(7);981–994. pmid:15946294

43. Kim J, Hahn JS, Franklin MJ, Stewart PS, Yoon J. Tolerance of dormant and active cells in Pseudomonas aeruginosa PAO1 biofilm to antimicrobial agents. J Antimicrob Chemother. 2009;63(1);129–135. pmid:19001451

44. De Groote VN, Verstraeten N, Fauvart M, Kint CI, Verbeeck AM, Beullens S, et al. Novel persistence genes in Pseudomonas aeruginosa identified by high-throughput screening. FEMS Microbiol Lett. 2009;297;73–79. pmid:19508279

45. Jubair M, Morris JG Jr, Ali A. Survival of Vibrio cholerae in nutrient-poor environments is associated with a novel “persister” phenotype. PLoS ONE. 2012;7(9); e45187. pmid:23028836

